# Complete persistence of the primary somatosensory system in zebrafish

**DOI:** 10.1101/2023.12.19.572352

**Authors:** Joaquín Navajas Acedo

## Abstract

The somatosensory system detects peripheral stimuli that are translated into behaviors necessary for survival. Fishes and amphibians possess two somatosensory systems in the trunk: the primary somatosensory system, formed by the Rohon-Beard neurons, and the secondary somatosensory system, formed by the neural crest cell-derived neurons of the Dorsal Root Ganglia. Rohon-Beard neurons have been characterized as a transient population that mostly disappears during the first days of life and is functionally replaced by the Dorsal Root Ganglia. Here, I comprehensively follow Rohon-Beard neurons *in vivo* and show that the entire repertoire remains present in zebrafish from 1-day post-fertilization until the juvenile stage, 15-days post-fertilization. These data indicate that zebrafish retain two complete somatosensory systems until at least up to a developmental stage when the animals display complex behavioral repertoires.

## Introduction

The somatosensory system of animals detects peripheral stimuli, such as touch, temperature, or noxious chemicals (Abraira and Ginty, 2013; Basbaum et al., 2009; Cevikbas and Lerner, 2020; Dhaka et al., 2006; Meltzer et al., 2021, 2021; Woolf and Ma, 2007). In vertebrates, the somatosensory system of the head is formed by neurons of the Trigeminal Ganglia (Dyck and Thomas, 2005). In the trunk, the situation varies between amniotes and anamniote vertebrates such as fishes and amphibians. While the somatosensory system of the trunk in amniotes is formed by neurons of the Dorsal Root Ganglia (DRG), which are neural crest cell-derived (Le Douarin and Kalcheim, 1999), anamniote vertebrates possess two somatosensory systems during development: the primary somatosensory system and the secondary somatosensory system. The primary somatosensory system develops first and is formed by the Rohon-Beard (RB) neurons (Beard, 1890; Bernhardt et al., 1990; Coghill, 1914; Freud, 1878, 1877; Hughes, 1957; Ogino and Hirata, 2018; Rohon, 1884). RB neurons are bipolar neurons present on the dorsal part of the spinal cord, and participate in the escape response (Clarke et al., 1984; Hartenstein, 1993; Hirata and Iida, 2018; Kimmel and Westerfield, 1990; Roberts and Clarke, 1982; Roberts and Smyth, 1974; Shorey et al., 2021; Umeda et al., 2016). RB neurons possess a characteristic large spherical body and extend their highly arborized sensory neurites to the periphery around 18 hours post-fertilization (hpf) in zebrafish (Eisen, 1991; Sagasti et al., 2005; Saint-Amant and Drapeau, 1998). Around the same stage, the neural crest cells of the trunk migrate out of the neural tube (Raible et al., 1992; Theveneau and Mayor, 2012) and start differentiating into DRGs, the secondary somatosensory system (An et al., 2002; Raible et al., 1992; Raible and Eisen, 1994; Wright and Ribera, 2010). Concomitantly to the maturation of the DRGs, RBs are thought to undergo gradual programmed cell death, and their replacement by the DRG is considered to be complete at around 5 days post-fertilization (dpf) in zebrafish (Cole and Ross, 2001; Lamborghini, 1987; Reyes et al., 2004; Svoboda et al., 2001; Williams et al., 2000). RBs show markers for cell death, including Terminal deoxynucleotidyl transferase dUTP nick end labeling (TUNEL), activated Caspase 3, or Annexin V during their disappearance and functional replacement by the DRGs in fish and frogs (Coen et al., 2001; Cole and Ross, 2001; Kanungo et al., 2006; Reyes et al., 2004; Svoboda et al., 2001, 2001; Williams et al., 2000; Williams and Ribera, 2020). While programmed cell death of RBs is dependent on electrical activity, neurotrophin, Cdk5 or BCL signaling (Coen et al., 2001; Kanungo et al., 2006; Nakano et al., 2010; Ogino and Hirata, 2018; Pineda et al., 2006; Svoboda et al., 2001; Williams et al., 2000), it is independent of the formation of DRGs (Honjo et al., 2011; Reyes et al., 2004).

RBs were originally regarded to completely disappear during development; however, recent observations in zebrafish challenged this view and documented that up to 40% of the RBs survive to juvenile stages (Palanca et al., 2013; Williams and Ribera, 2020). Here, I report that the complete repertoire of RBs present at 1 dpf remains until 15 dpf and show no significant signs of cell death.

## Results

### The complete trunk repertoire of RBs survives until at least 5 dpf in zebrafish

To understand the development of the primary somatosensory system through the dynamics of RBs loss, I investigated *(i)* where in the trunk and *(ii)* when each of the RBs disappears. I repeatedly and comprehensively followed *in vivo* the RBs of individual fish daily from 1 dpf until 5 dpf using an *isl2b:GFP* transgenic line that labels all RBs (Palanca et al., 2013; Pittman et al., 2008) (Figure 1). The *isl2b:GFP* transgenic line labels both RBs and Dorsal Longitudinal Ascending (DoLA) interneurons throughout early development, and additionally DRG neurons from 2-3 dpf onwards (Williams and Ribera, 2020; Won et al., 2012). RBs were identified by their dorsal position and characteristic large soma, while DoLA were identified based on their smaller size and adjacent position to the dorsal longitudinal fasciculus (Figure 1). DRGs were identified based on their location outside the spinal cord. RBs start as two bilateral populations that converge medially (Fig. 1, compare top vs bottom panel) (Williams and Ribera, 2020). Despite their change in position, in all embryos analyzed (n=3), all trunk *isl2b:GFP+* RB neurons present at 24 hpf could be accounted for throughout the entirety of the experimental period, until 5 dpf. These results indicate that all RBs survive past hatching (∼2-3 dpf) until free-swimming larva stages (5 dpf).

**Figure 1.**
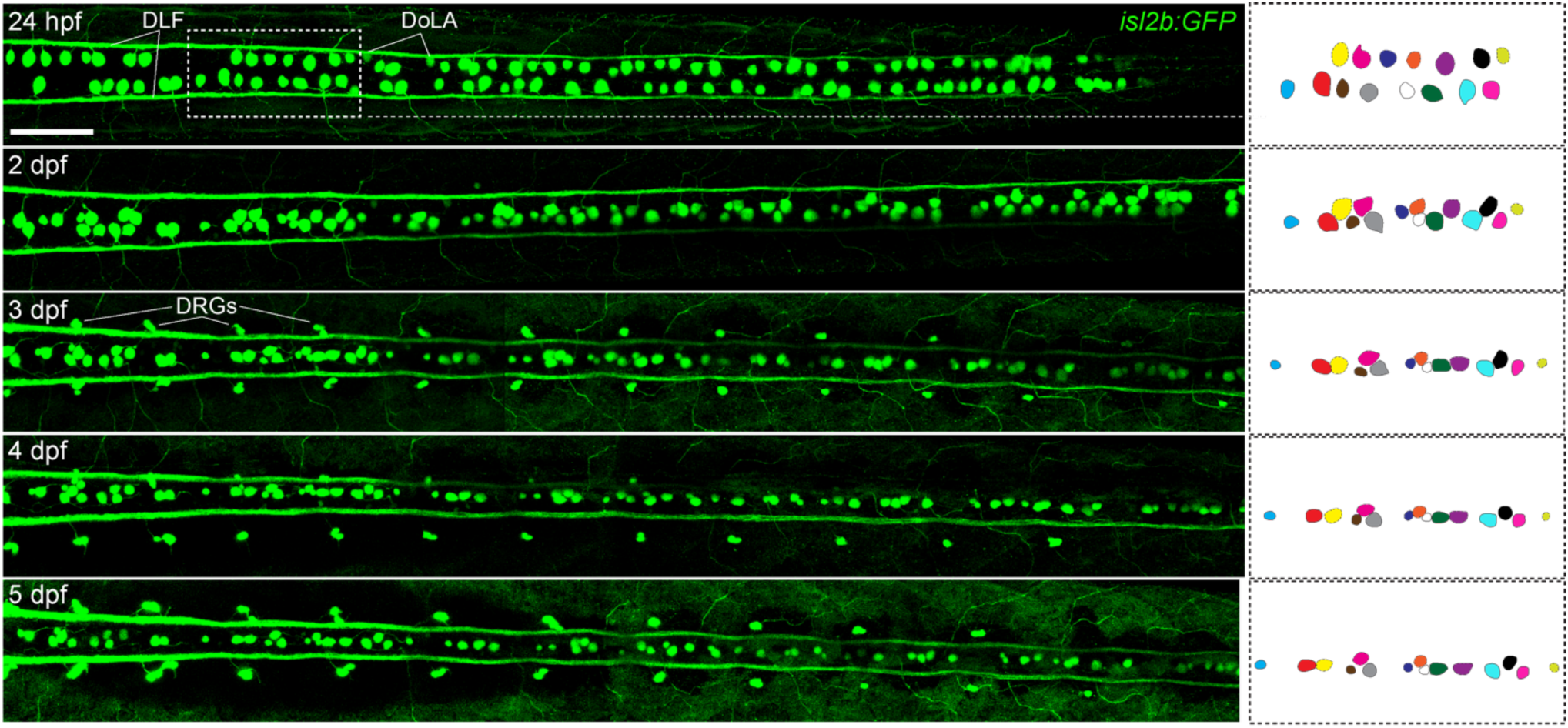
All *isl2b:GFP+* RBs present at 24 hpf can be accounted for at 5 dpf. Repeated dorsal *in vivo* imaging of the same larva from the first until the fifth day of life reveals that all *isl2b:GFP+* RBs present in the trunk survive during this period of time. The drawings on the right represent the same *isl2b:GFP+* RBs throughout time, starting from the dotted box area at 24 hpf. Scale bar equals 100 µm. hpf – hours post-fertilization; dpf - days post-fertilization; DoLA – Dorsal Longitudinal Ascending interneurons; DRG – Dorsal Root Ganglia; DLF – Dorsal Longitudinal Fasciculi.

### The zebrafish trunk elongates as RB converge towards the midline

The comprehensive tracking revealed that all RBs are present until 5 dpf and their bodies converge medially, ending up in approximately a single row (Fig. 1)(Williams and Ribera, 2020). Interestingly, the bodies of *isl2b:GFP+* RB neurons seem to get displaced anterio-posteriorly (Fig. S1, compare top vs bottom panels). A time-lapse video of an *isl2b:GFP* transgenic line shows this effect *in vivo* from 1 to 2 dpf (Video 1). I then wondered if this displacement was exclusive to RBs or whether the surrounding tissues also participate. To characterize the ongoing elongation of the trunk and RB convergence to the midline, I injected *isl2b:GFP* embryos with mRNA encoding a photoconvertible nuclear-localized Kaede (Figure 2) and followed the photoconverted area and the individual *isl2b:GFP+* RBs. In all analyzed samples, the space between the two photoconverted areas increased between 24 hpf and 3 dpf (Fig. 2 top vs bottom; n=3, mean distance 1 vs 1.391, p-val= 0.0004). Together, these data indicate that there is not only a convergence of RBs towards the midline, but also anterio-posterior displacement due to the concomitant extension of the trunk.

**Figure 2.**
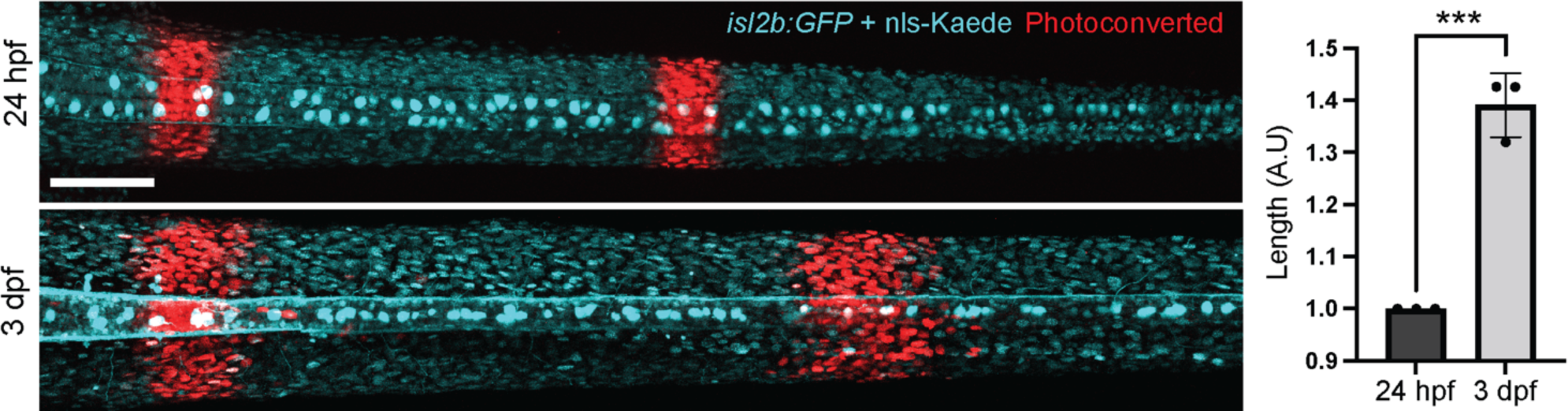
Photoconversion indicates the concomitant trunk elongation and medial convergence of RBs. Two stripes were photoconverted on the trunk of *isl2b:GFP* fish injected with a nuclear-localized Kaede (nls-Kaede). The images were aligned using the anteriormost photoconverted area, revealing the space between the two stripes increased (n=3; Mean length of ‘1’ at 24hpf, versus ‘1.391’ at 3 dpf). Asterisks indicate statistical significance. Scale bar equals 100 µm. hpf — hours post-fertilization; dpf — days post-fertilization.

**Video 1.**
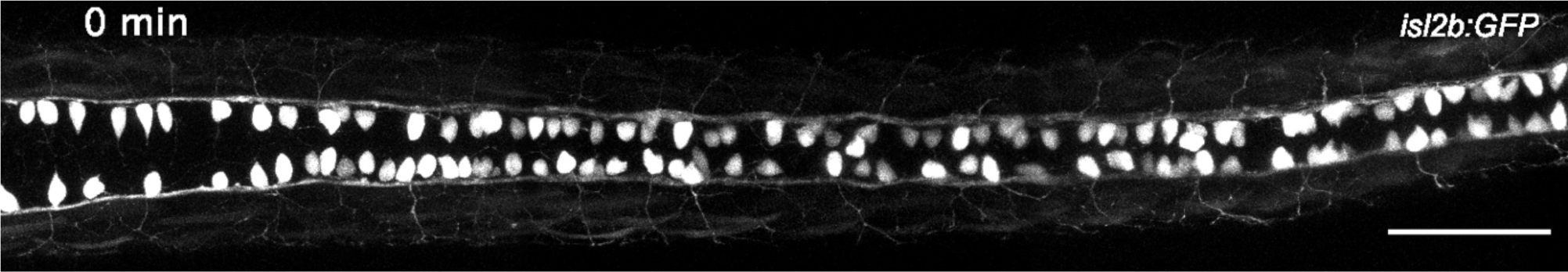
Time lapse of medial convergence of *isl2b:GFP+* RB neurons from 24 hpf to 48 hpf. *In vivo* time-lapse imaging of the dorsal spinal cord of a *isl2b:GFP* transgenic line from 24 to 48 hpf (Dorsal view). One frame corresponds to 30 minutes. Scale bar equals 100 µm.

### The vast majority of RBs survive until juvenile stages

The data above shows no decline in the number of RBs during the first days of life and full survival until 5 dpf (Fig. 1). To test how many *isl2b:GFP+* RBs survive to juvenile stages, I followed the same animals from 3 dpf —when the medial convergence has ended (Fig. 1)- until 15 dpf. *isl2b:GFP+* RBs can be identified at 15 dpf based on their medial location and expression of *isl2b:GFP* (Figure 3)(Won et al., 2012, 2011). As the animal grows in length, the space between *isl2b:GFP+* RBs continues to increase overall (compare Fig. 3 3 dpf vs 15 dpf). Furthermore, 97-100% of *isl2b:GFP+* cells – including RBs- present at 3 dpf survived until 15 dpf in my imaging conditions and analyzed animals (Fig. 3, Supplementary Table 1, n=5). Altogether, this data indicates that the vast majority of RBs survive until juvenile stages in zebrafish.

**Figure 3.**
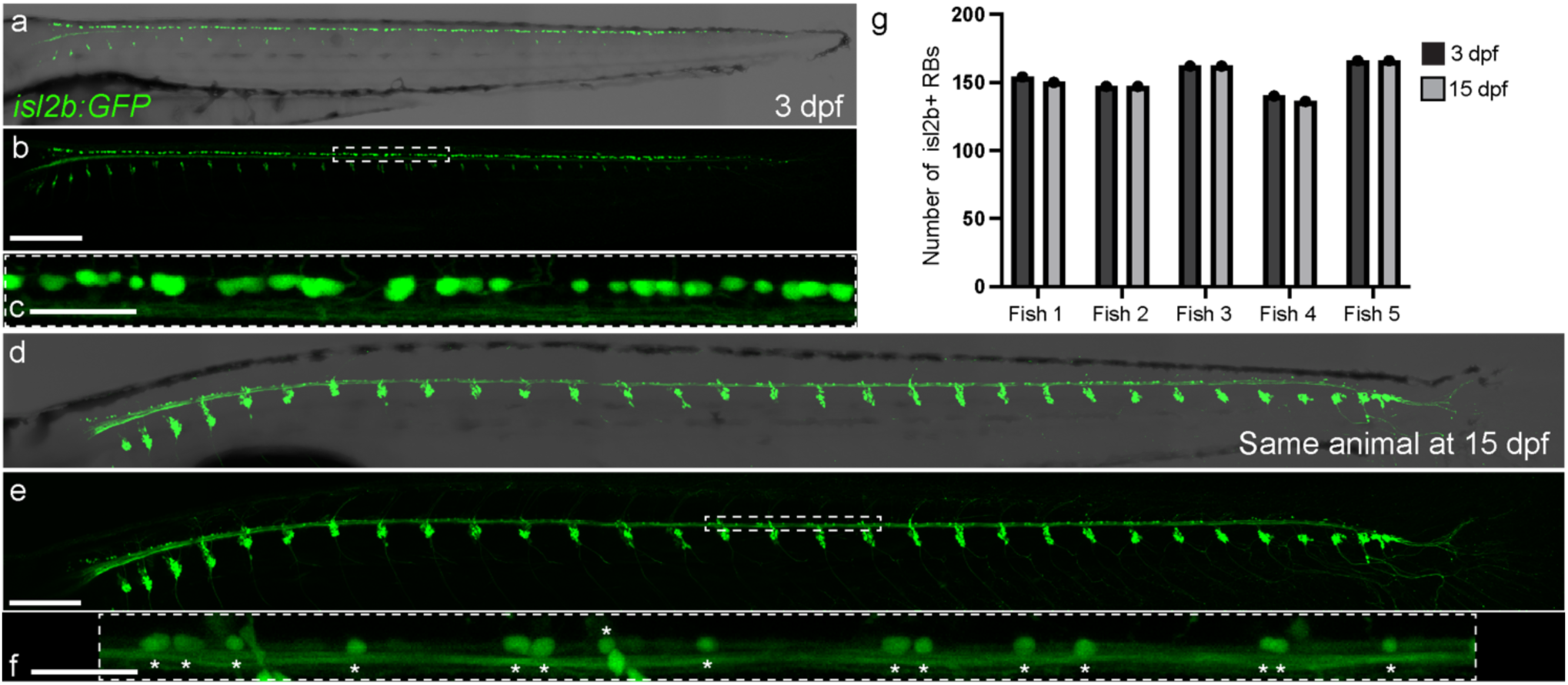
The vast majority of RBs survive until juvenile stages. Imaging of the same *isl2b:GFP* transgenic animal at 3 (a-c) and 15 dpf (d-f) show that the majority of *isl2b:GFP+* RBs are still present in the spinal cord of 15 dpf zebrafish and the distance between them increased. (c and f) Both images show the same area between DRGs number 16 and 19 (delineated in b and e). Asterisks in (f) label *isl2b:GFP+* RBs at 15 dpf. Because of the time resolution, the exact RB identity between the 3 and 15 dpf time points could not be established. (g) Quantification of the number of *isl2b:GFP+* RBs per animal at 3 and 15 dpf (exact numbers in Supplementary Table 1). Scale bars in (a-b and d-e) equal 250 µm, and 50 µm in (c) and (f). Images from 3 dpf and 15 dpf are to scale to each other to reflect the amount of growth. dpf — days post-fertilization.

### RBs do not show signs of programmed cell death at 24 hpf

Given the persistence of RBs through 5dpf, and until juvenile stages (Fig. 1 and 3, Video 1), I next tested whether cell death markers were distributed in any discernible pattern that may suggest a biological role other than cell death. To detect cell death markers, I used two different methods previously reported in RBs: a Sec5A-YFP fluorescent reporter, which labels fluorescently flipped phosphatidylserine groups in cells undergoing apoptosis (Ham et al., 2010; Williams and Ribera, 2020), and TUNEL (Reyes et al., 2004; Svoboda et al., 2001; Williams et al., 2000). In my assays, RBs did not show sec5A-YFP signal at 24 hpf (Figure 3a, Fig. S2, n=12); however, other previously reported cells types outside the spinal cord using this transgenic line showed reproducible secA5-YFP signal, confirming the functionality of the reporter in my experimental setting (Fig. 3a, arrowheads)(Ham et al., 2010). Furthermore, RBs did not show significant TUNEL staining (Figure 3b-d). In all animals analyzed (n=9), only one TUNEL+ RB neuron was found, but other nearby cells in the skin and spinal cord did show robust TUNEL staining, confirming the functionality of the assay (Fig. 3b-c). Together with my *in vivo* imaging and comprehensive tracking, these results argue for a persistence of RB neurons in zebrafish until at least 15 dpf with little to no reduction of initial RB neuron numbers due to programmed cell death.

## Discussion

Since their first description in the late 1800s (Beard, 1890; Freud, 1878, 1878; Rohon, 1884), RB neurons have attracted the attention of both developmental biologists and neuroscientists due to their large soma size, their accessibility to electrophysiological recordings and imaging, and their function in somatosensation and escape response (Artinger et al., 1999; Bernhardt et al., 1990; Blader et al., 2003; Douglass et al., 2008; Henderson et al., 2020, 2019; Hubbard et al., 2016; Jacobson, 1981; Kaji and Artinger, 2004; Knafo et al., 2017; Lamborghini, 1987; Moreno and Ribera, 2014; Nieuwenhuys, 1964; O’Brien et al., 2012; Ogino and Hirata, 2018; Park et al., 2012; Rossi et al., 2009, 2008; Spitzer, 1984, 1982). Previous reports described the total or partial disappearance of RB neurons starting at around the first day of life in fishes (Bernhardt et al., 1990; Henion et al., 1996; Metcalfe et al., 1990; Metcalfe and Westerfield, 1990; Ogino and Hirata, 2018; Reyes et al., 2004; Svoboda et al., 2001; Williams et al., 2000; Williams and Ribera, 2020). In contrast to these observations, the data presented here indicate that the entire repertoire of RB neurons survives until juvenile stages. First, comprehensive daily tracking between 1 and 5 dpf shows that all RBs can be accounted for even if they undergo medial and anterio-posterior displacement. Second, the vast majority if not all RBs survive until 15 dpf. Finally, RBs do not display a significant presence of classical apoptotic markers.

### Revealing the persistence of RB neurons with live imaging

What factors and experimental settings might have led to the conclusion that RB neurons disappear during zebrafish development? Previous characterizations of RB disappearance during development were based on either *(i)* reporting an average number of RBs per somite, *(ii)* imaging of animals laterally rather than dorsally, *(iii)* quantification of RB number in fixed animals, and/or *(iv)* use of markers (e.g HNK1/zn-12, *isl1SS* enhancer) that might stop being expressed or not be expressed in all RBs (Appel et al., 1995; Eisen and Pike, 1991; Grunwald et al., 1988; Harris and Whiting, 1954; Joya et al., 2014; Metcalfe et al., 1990; Nakano et al., 2010; Nordlander, 1989; Palanca et al., 2013; Pineda et al., 2006; Reyes et al., 2004; Takamiya and Campos-Ortega, 2006; Tamme et al., 2002; Tongiorgi et al., 1995; Uemura et al., 2005; Williams et al., 2000; Williams and Ribera, 2020; Won et al., 2012, 2011). Furthermore, recent work demonstrated that the characteristic large RB soma size decreases over time (Williams and Ribera, 2020), making it difficult to differentiate RBs from other spinal cord neurons using antibodies such as Isl1/2/39.4D5 or Elavl3/HuC (Rossi et al., 2009; Segawa et al., 2001).

Considering the comprehensive *in vivo* tracking data presented here (Fig. 1), the fact that RBs converge medially and that the trunk extends concomitantly (Fig. 2 and 3, and Video 1), it is possible that the combination of these processes has contributed to the interpretation of RB numbers decreasing over time. For example, fixed samples of different animals or less comprehensive tracking in time might suggest that RBs disappear when in fact they redistribute along the elongating trunk and thus *de facto* decrease their density. The comprehensive longitudinal tracking method employed in this work complements prior approaches, and underlines the utility of tracking all RBs on a per-animal basis in the context of concurrent developmental processes. Cell death by apoptosis critically contributes to sculpting the nervous system during development (Burek and Oppenheim, 1996; Charvet et al., 2011; Dekkers et al., 2013; Malin and Shaham, 2015; Pop et al., 2020). However, in my described assays and conditions, I did not detect any significant presence of cell death markers in RB neurons (Fig. 4). Previous publications have described the presence of cell death markers in RB neurons, including activated Caspase-3, TUNEL and Annexin V (Coen et al., 2001; Cole and Ross, 2001; Ham et al., 2010; Kanungo et al., 2006; Reyes et al., 2004; Svoboda et al., 2001, 2001; Williams et al., 2000; Williams and Ribera, 2020). These reports showed that not all cells that are secA5-YFP+ are TUNEL+ (Dong et al., 2011; Ham et al., 2010; Williams and Ribera, 2020), and a negligible number of RBs express activated Caspase-3 per embryo (Williams and Ribera, 2020). These observations contrast with other populations of neurons that undergo apoptosis in zebrafish (Mazaheri et al., 2014). While apoptosis of a few cells cannot be ruled out, my observations using SecA5-YFP and TUNEL indicate that cell death is not a common fate of RB neurons in the first 15 days of zebrafish development.

**Figure 4.**
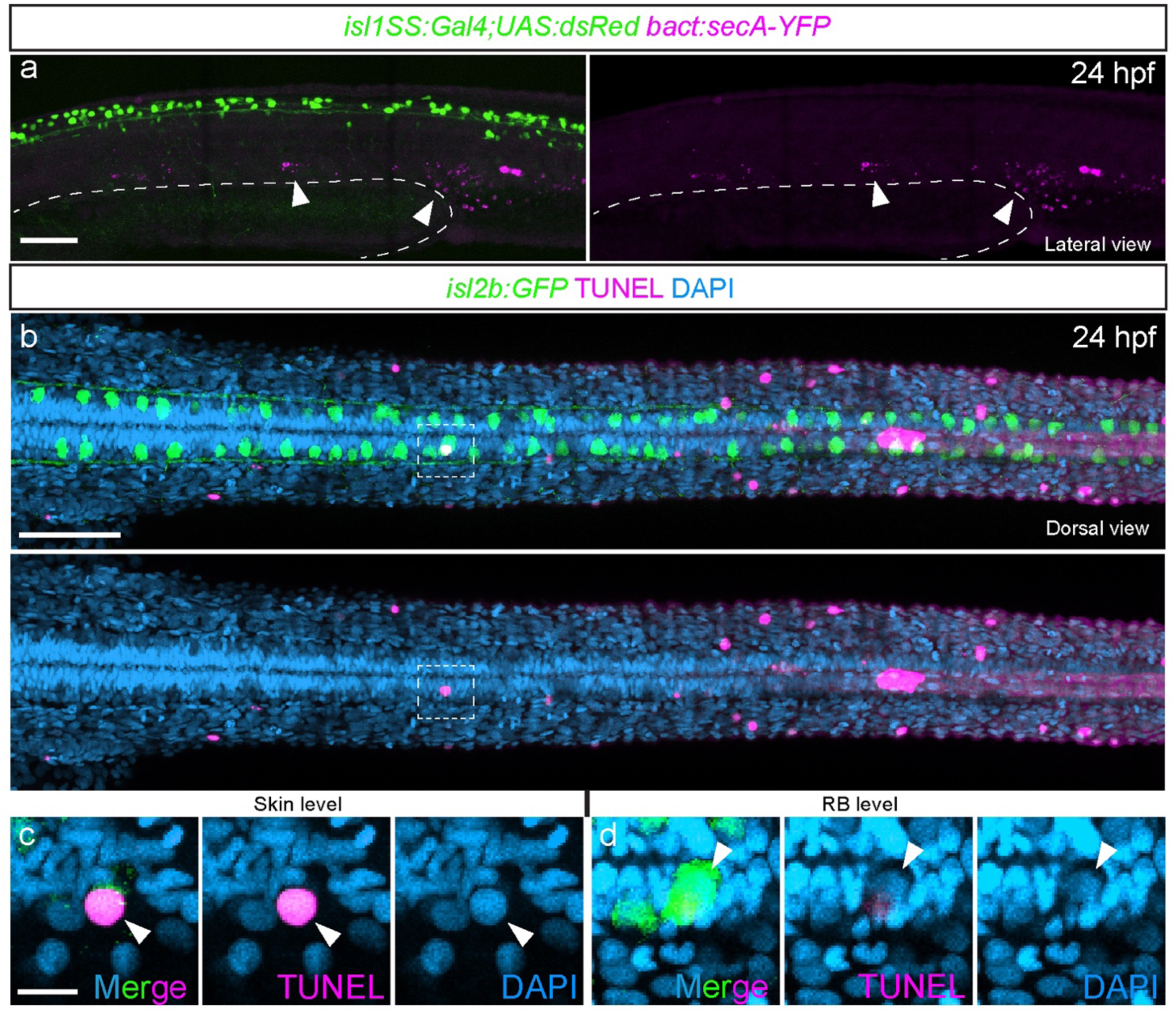
RBs do not show SecA5-YFP or TUNEL signal at 24hpf. Images showing lateral (upper; a) and dorsal views (lower; b-d) of two different experiments assessing cell death. (a) secA5-YFP is driven by a ubiquitous beta-Actin promoter, and none of the dsRed+ RBs were secA-YFP+. Some cells around the pronephros area were were SecA5-YFP+ (arrowheads) (n=12). The yolk extension is delineated by a dashed line. (b) A dorsal view of an *isl2b:GFP+* embryo stained for TUNEL. None of the RBs were TUNEL positive, but some surrounding cells were (n=9). Inset (dashed line in b) shows the TUNEL+ nucleus of a skin cell (c, arrowhead) and the RB immediately underneath (d, arrowhead). Scale bars equal 100 µm in a-b, and 10 µm in c-d, respectively. hpf — hours post-fertilization.

Independently from the work in this manuscript, Liu and Kucenas (Liu and Kucenas 2024, *under review*) reached similar conclusions: RB neurons do not disappear during early zebrafish development and do not show canonical markers for programmed cell death. Liu and Kucenas used a different set of transgenes that are expressed in RBs (−3.1*neurog1:GFP^sb2^*, *neurod1:Gal4^uva22^*)(Blader et al., 2003; Fontenas and Kucenas, 2021) to track their presence across development. The authors found that RB neurons persist until at least 16 dpf, in agreement with the data in this manuscript. Furthermore, the authors did not find any significant presence of similar programmed cell death markers than the ones used in this manuscript (TUNEL, secA5-YFP) in RBs, providing further evidence RB neurons persist during development of zebrafish and are not eliminated by using that mechanism.

If cell death markers are not necessarily labeling programmed cell death in RBs, what concomitant process might they reflect? The cell death machinery has non-apoptotic roles in neurons and is involved in axon pruning or pathfinding (Mukherjee and Williams, 2017). Furthermore, double strand breaks in the DNA contribute to neuron maturation, through control of gene diversification, induction of gene expression, or cytoskeletal dynamics (Akagawa et al., 2021; Alt and Schwer, 2018; Álvarez-Lindo et al., 2022; Kellermeyer et al., 2018; Madabhushi et al., 2015). RB neurons expressing markers associated with cell death might instead still be in the process of maturation or remodeling their sensory axons while reaching their peripheral targets. Future work will shed light on the role that the cell death machinery may have during development of RBs.

### Outlook

The data presented here show that the entire repertoire of zebrafish RB neurons that are present during the first day of life persists until juvenile stages. My study raises several questions about the fate and roles of RBs. First, is survival of RBs exclusive of fishes? The data presented here argues against prominent neuron disappearance through programmed cell death during development of the somatosensory system of zebrafish. However, this situation might be exclusive to fishes and not translate to amphibian models, which possess RBs during early stages of development but undergo a more dramatic metamorphosis process that involves major remodeling of the nervous system (Coen et al., 2001; Coghill, 1914; Eichler and Porter, 1981; Kerr et al., 1972; Kollros and Bovbjerg, 1997; Lamborghini, 1987; Nishikawa, 2012).

Second, do RBs act as pioneer neurons upon which the DRG system builds? RBs and DRGs overlap during a substantial amount of time (Fig. 1 and 3), including the period of free-swimming larva and acquisition of complex behaviors. Several systems are built using pioneer neurons, including innervation of the limbs in grasshoppers (Bentley and Keshishian, 1982), CNS development in Drosophila (Hidalgo and Brand, 1997), Cajal-Retzius and subplate neurons in mammals (Frotscher, 1997; McConnell et al., 1989; Meyer et al., 1998), and neurons of the CNS, statoacoustic ganglion and olfactory system in zebrafish (Bañón and Alsina, 2023; Whitlock and Westerfield, 1998; Wilson and Easter, 1991). RB and DRG peripheral arbors overlap (Wright and Ribera, 2010), and while the DRGs are not necessary for RB development (Honjo et al., 2011; Reyes et al., 2004), whether RB neurons participate during the development of the secondary somatosensory system is unknown.

Third, what are the physiological and behavioral consequences of having two functional somatosensory systems in the trunk? Do RBs and DRG neurons complement each other functionally? In zebrafish, DRGs and RBs express a shared set of receptors and signaling molecules related to their somatosensory function, including NTRKs, purinergic receptors, *scn8aa*/Nav1.6, and PKCa (Kucenas et al., 2006; Martin et al., 1998; Palanca et al., 2013; Patten et al., 2007; Pineda et al., 2006; Tuttle et al., 2024; Won et al., 2012). Both RBs and DRGs innervate the skin and specialized structures such as the pectoral fin and scales (Henderson et al., 2020, 2019; O’Brien et al., 2012; Rasmussen et al., 2018), and their sensory processes overlap substantially (Wright and Ribera, 2010). Future studies comparing and manipulating the activity and function of RBs and DRGs will help reveal the overlapping or split roles these somatosensory systems may have. The work presented here, and by Liu and Kucenas, provide a new, exciting outlook in the field of RB and somatosensory system development.

## Materials and Methods

### Fish Husbandry

Zebrafish lines were raised under standard light-dark conditions (14-10 hours) and fed a standard diet of Artemia and Dry food 4 times a day. All zebrafish experiments were performed according to the Swiss Law and the Kantonales Veterinäramt of Kanton Basel-Stadt (licenses #1035H and #3097). Wild-type TLAB, *Tg(−17.6kb isl2:mmGFP5)^cz7^* (referred to as *isl2b:GFP* throughout the manuscript, (Pittman et al., 2008); a kind gift from Dr. Berta Alsina), *Tg(isl1SS:Gal4;UAS:dsRed)^zf234Tg^* (a kind gift from Dr. A Sagasti; *sensory:RFP* in Palanca et al. (Palanca et al., 2013)) and *Tg(bAct:secA-YFP)* (this manuscript and (Ham et al., 2010)) zebrafish lines were used. Embryos were obtained by standard cross protocol, and collected in E3 embryo media (5 mM NaCl, 0.17 mM KCl, 0.33 mM CaCl2, 0.33 mM MgSO4, pH 7.2) and kept in an incubator at 28.5C and a 14h light/10h dark cycle. For the experiments that required from repeated imaging from 24 hpf to 5 dpf, 0.003% 1-phenyl 2-thiourea (PTU) was added to the embryo media to block the formation of pigment. Staging of the embryos was performed according to Kimmel et al. (Kimmel et al., 1995), available at the Zebrafish Information Network (ZFIN) (Bradford et al., 2022).

### Tg(bAct:secA-YFP) transgenic fish

To generate the *Tg(bAct:secA-YFP) ^a400Tg^* transgenic fish, the Tol2-bActin:secA-YFP plasmid (Addgene #105664, a gift from Dr. Qing Deng) was co injected with Tol2 transposase mRNA as previously described (Suster et al., 2009) into one-cell stage embryos. Then, embryos were screened at 24 hpf for positive YFP clones and raised until adulthood. The F0 founders were screened for germline integration of *bAct:secA-YFP* by outcrossing to TLAB wild-type fish, and the F1 progeny raised until adulthood. Afterwards, F1 stable fish were screened for robust YFP expression by outcross to TLAB wild-type fish, and the produced F2 progeny raised until adulthood. All experiments were conducted using F3s of the generated *Tg(bAct:secA-YFP)^a400Tg^* transgenic lines.

### Zebrafish mounting and time-lapse Imaging

*isl2b:GFP* positive fish were manually dechorionated, anesthetized using 6.5% MS-222/Tricaine Methanesulfonate (Sigma-Aldrich, Cat#E10521; 4g/L pH9.0 stock in E3 embryo media) and mounted in Low Melting Point Agarose (LMP; Sigma-Aldrich, Cat#A-9414) as previously described (Venero Galanternik et al., 2016) in Glass-Bottom MatTek plates (MatTek). For dorsal views, the dorsal part of the animal was closer to the coverslip, and fluorescent images were acquired using a Zeiss LSM880 AiryScan, 25X Oil Objective in a heated chamber at 28.5C. The produced time lapse video was time registered using the ‘*stackreg*’ plugin (https://research.stowers.org/imagejplugins/ImageJ_tutorial2.html) to maintain the original center of the field of view stable.

For the time-lapses that required from repeated imaging every 24 hours in Figure 1, after each session of imaging (approximately 30-45 minutes long), larvae were retrieved from the LMP and kept in fresh E3 embryo media + 0.003% PTU until the next imaging session the next day. Images were processed with the commercial software Adobe Photoshop 2021 for Intensity and Contrast, and the Figures were assembled in Adobe Illustrator 2021.

### Photoconversion

mRNA was synthetized using the SP6 mMessenger Machine (Thermo Fisher, Cat#AM1340) and purified using Zymo RNA Cleaner and Concentrator (Zymo Research, Cat#R1017). nlsKaede mRNA (a Kind gift from Dr. K Kwan (Kwan et al., 2012)) was injected into one-cell stage eggs from an *isl2b:GFP* to TLAB cross, under standard conditions. The mRNA was injected at a concentration of 20 pg per embryo, and eggs were kept in an incubator at 28.5C in the dark to avoid fluorophore bleaching. At 23 hpf embryos were screened for robust expression of both nls-Kaede and the *isl2b:GFP* transgene, and mounted for imaging in LMP as described above.

Photoconversion was performed using the 405 UV laser of a Zeiss LSM880 AiryScan, 25X Oil Objective, using the ‘Regions’ and ‘Bleaching Setup’ of ZEN Software until photoconverted nuclear Kaede intensity was robust.

### Statistical analysis

Distances between photoconverted areas in Figure 2 were measured in Fiji (Schindelin et al., 2012) using the ‘Line’ tool, and converted to ratios of 24 hpf vs 72 hpf values. Unpaired t-test statistical analysis and data plotting were performed using the commercial software Prism 9.

### Terminal deoxynucleotidyl transferase dUTP nick end (TUNEL) staining

*isl2b:GFP* positive larvae were fixed using 4% PFA in 1X PBS (pH=7.4) at 24 hpf. TUNEL staining was performed using the In Situ Cell Death Detection Kit (Roche, Cat #11684817910) according to the manufacturer’s specifications. Anti-DIG (in the kit) and anti-GFP (Biozol, Aves GFP-1020; 1:500) antibodies were co-incubated.

## Acknowledgements

I thank Alex Schier for early input and enthusiastic support of this work, the members of the Schier Lab for critical feedback on the project, and Mariona Colomer Rosell, Mireia Codina Tobias, Drs. Jialin Liu, Lucia Du, Oded Mayseless, Clare Diester, Madalena Pinto, Yinan Wan, and Alex Schier for critical reading of the manuscript. Thanks to Drs. Christian Mosimann, Catherina Becker, Dave Raible, Patrick Blader, and Jeff Rasmussen for comments on the manuscript and discussion of the data. I additionally thank Alba Aparicio Fernandez, Rita Gonzalez and Diana Medeiros Gomes for outstanding zebrafish husbandry, and the Biozentrum’s Imaging Core Facility (IMCF) for their infrastructure support and assistance. I thank Dr. Berta Alsina for sending the *isl2b:GFP* fish line, Dr. Kristen Kwan for the pCS2-nls-Kaede plasmid, and Drs. Christian Mosimann, Tim Saunders and Kostantinos Kalyviotis for providing reagents during pilot experiments. I would like to thank Kendra Liu and Dr. Sarah Kucenas for our scientific exchanges and discussions during the process of co-submission. Many thanks to ZFIN (www.zfin.org) for being an unvaluable resource for this project and the entirety of the zebrafish community. The project was supported by the Allen Discovery Center for Cell Lineage Tracing.

## Declaration of generative AI in scientific writing

I declare no generative AI was used at any stage during the writing of the manuscript

## Declaration of sex- and gender-based analyses

Sex determination in zebrafish occurs after juvenile stages, and no sex-determining genes have been described in zebrafish lines kept in the laboratory. I do not anticipate sex-biases to affect t the main conclusions of this work. Nevertheless, the somatosensory system is present in males and females, and the findings from this work can be applied to both sexes.

## Supplementary Figures

**Supplementary Figure 1 to Figure 1.**
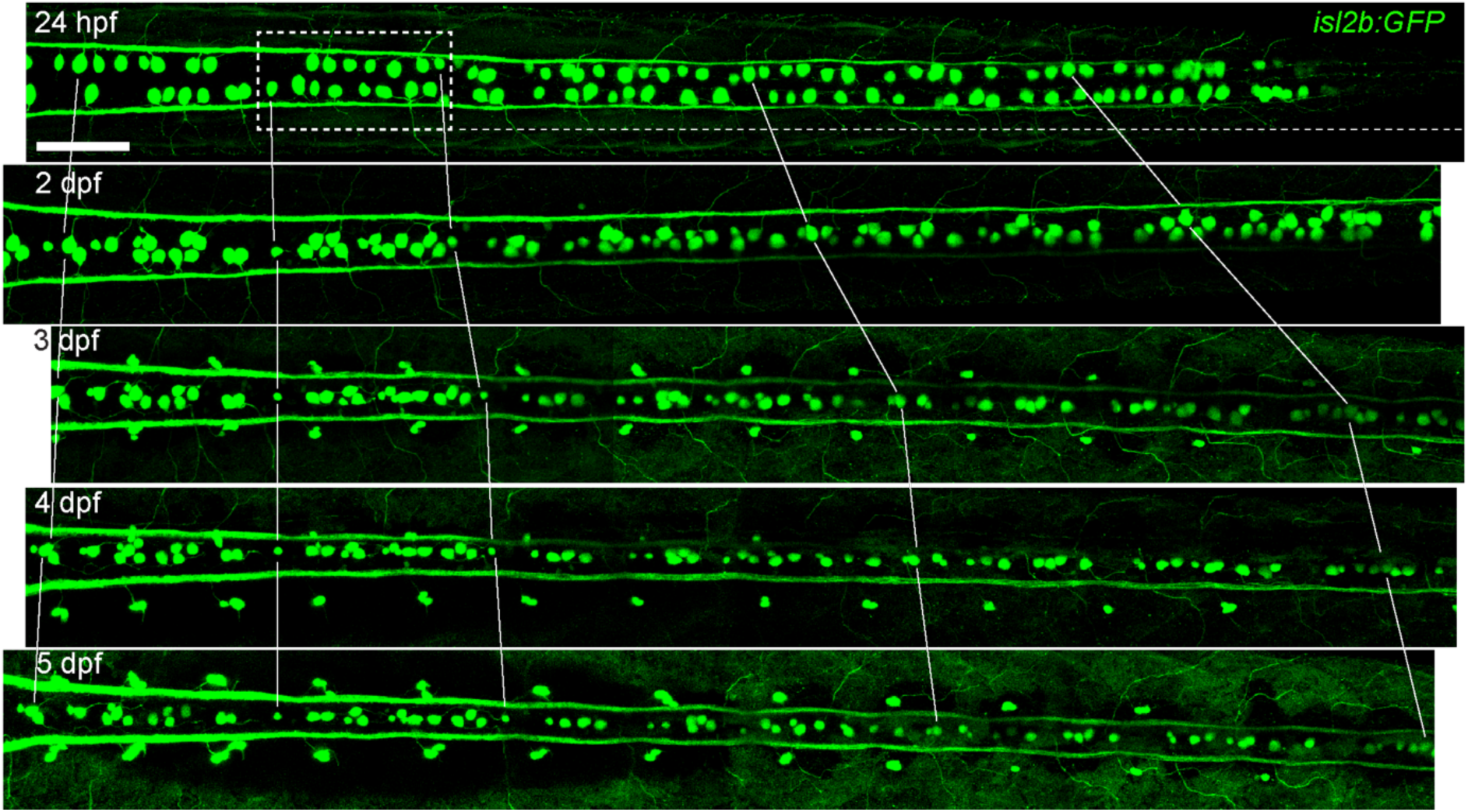
Lines show the amount of displacement the *isl2b:GFP+* RBs undergo in the anterio-posterior axis in Figure 1. Scale bar equals 100 µm. hpf — hours post-fertilization; dpf — days post-fertilization.

**Supplementary Figure 2 to Figure 4.**
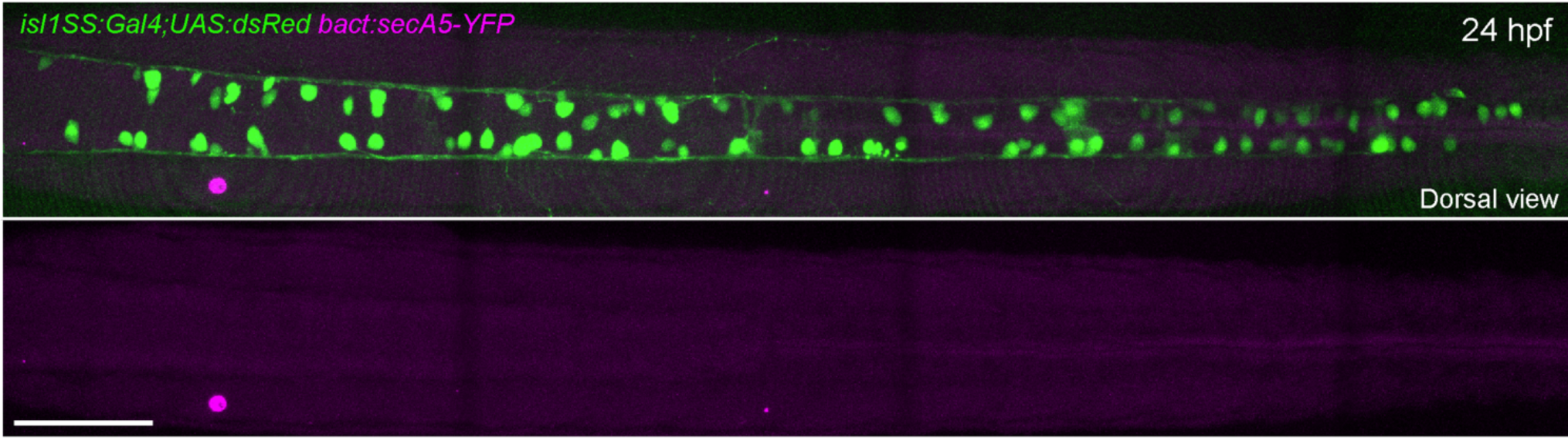
RBs do not show SecA5-YFP (Annexin V) signal at 24 hpf. Images showing a dorsal view of a *secA5-YFP+* transgenic embryo. *secA5-YFP* is driven by a ubiquitous beta-Actin promoter. None of the RBs of the trunk were *secA5-YFP+*. Scale bar equals 100 µm. hpf — hours post-fertilization.

**Supplementary Table 1 to Figure 3.** *isl2b:GFP+* RB counts of the same animals at 3 and 15 dpf (Fish 5 is shown in Figure 3).

| Age | # of *isl2b:GFP*<sup>+</sup> RBs |  |
| --- | --- | --- |
|  | 3 dpf | 15 dpf |
| Fish 1 | 154 | 150 |
| Fish 2 | 147 | 147 |
| Fish 3 | 162 | 162 |
| Fish 4 | 140 | 136 |
| Fish 5 | 166 | 166 |

## References

Abraira, V.E., Ginty, D.D., 2013. The Sensory Neurons of Touch. Neuron 79, 618–639. 10.1016/j.neuron.2013.07.051

Akagawa, R., Nabeshima, Y., Kawauchi, T., 2021. Alternative Functions of Cell Cycle-Related and DNA Repair Proteins in Post-mitotic Neurons. Front. Cell Dev. Biol. 9, 753175. 10.3389/fcell.2021.753175

Alt, F.W., Schwer, B., 2018. DNA double-strand breaks as drivers of neural genomic change, function, and disease. DNA Repair 71, 158–163. 10.1016/j.dnarep.2018.08.019

Álvarez-Lindo, N., Suárez, T., De La Rosa, E.J., 2022. Exploring the Origin and Physiological Significance of DNA Double Strand Breaks in the Developing Neuroretina. IJMS 23, 6449. 10.3390/ijms23126449

An, M., Luo, R., Henion, P.D., 2002. Differentiation and maturation of zebrafish dorsal root and sympathetic ganglion neurons. J of Comparative Neurology 446, 267–275. 10.1002/cne.10214

Appel, B., Korzh, V., Glasgow, E., Thor, S., Edlund, T., Dawid, I.B., Eisen, J.S., 1995. Motoneuron fate specification revealed by patterned LIM homeobox gene expression in embryonic zebrafish. Development 121, 4117–4125. 10.1242/dev.121.12.4117

Artinger, K.B., Chitnis, A.B., Mercola, M., Driever, W., 1999. Zebrafish narrowminded suggests a genetic link between formation of neural crest and primary sensory neurons. Development 126, 3969–3979. 10.1242/dev.126.18.3969

Bañón, A., Alsina, B., 2023. Role of pioneer neurons and neuroblast behaviors on otic ganglion assembly (preprint). Developmental Biology. 10.1101/2023.03.30.534903

Basbaum, A.I., Bautista, D.M., Scherrer, G., Julius, D., 2009. Cellular and Molecular Mechanisms of Pain. Cell 139, 267–284. 10.1016/j.cell.2009.09.028

Beard, J., 1890. IV. On the early development of Lepidosteus osseus . — Preliminary notice. Proc. R. Soc. Lond. 46, 108–118. 10.1098/rspl.1889.0015

Bentley, D., Keshishian, H., 1982. Pathfinding by Peripheral Pioneer Neurons in Grasshoppers. Science 218, 1082–1088. 10.1126/science.218.4577.1082

Bernhardt, R.R., Chitnis, A.B., Lindamer, L., Kuwada, J.Y., 1990. Identification of spinal neurons in the embryonic and larval zebrafish. J of Comparative Neurology 302, 603–616. 10.1002/cne.903020315

Blader, P., Plessy, C., Strähle, U., 2003. Multiple regulatory elements with spatially and temporally distinct activities control neurogenin1 expression in primary neurons of the zebrafish embryo. Mechanisms of Development 120, 211–218. 10.1016/S0925-4773(02)00413-6

Bradford, Y.M., Van Slyke, C.E., Ruzicka, L., Singer, A., Eagle, A., Fashena, D., Howe, D.G., Frazer, K., Martin, R., Paddock, H., Pich, C., Ramachandran, S., Westerfield, M., 2022. Zebrafish information network, the knowledgebase for *Danio rerio* research. Genetics 220, iyac016. 10.1093/genetics/iyac016

Burek, M.J., Oppenheim, R.W., 1996. Programmed Cell Death in the Developing Nervous System. Brain Pathology 6, 427–446. 10.1111/j.1750-3639.1996.tb00874.x

Cevikbas, F., Lerner, E.A., 2020. Physiology and Pathophysiology of Itch. Physiological Reviews 100, 945–982. 10.1152/physrev.00017.2019

Charvet, C.J., Striedter, G.F., Finlay, B.L., 2011. Evo-Devo and Brain Scaling: Candidate Developmental Mechanisms for Variation and Constancy in Vertebrate Brain Evolution. Brain Behav Evol 78, 248–257. 10.1159/000329851

Clarke, J.D., Hayes, B.P., Hunt, S.P., Roberts, A., 1984. Sensory physiology, anatomy and immunohistochemistry of Rohon-Beard neurones in embryos of Xenopus laevis. J Physiol 348, 511–525. 10.1113/jphysiol.1984.sp015122

Coen, L., du Pasquier, D., Le Mevel, S., Brown, S., Tata, J., Mazabraud, A., Demeneix, B.A., 2001. Xenopus Bcl-XL selectively protects Rohon-Beard neurons from metamorphic degeneration. Proceedings of the National Academy of Sciences 98, 7869–7874. 10.1073/pnas.141226798

Coghill, G.E., 1914. Correlated anatomical and physiological studies of the growth of the nervous system of amphibia. J of Comparative Neurology 24, 161–232. 10.1002/cne.900240205

Cole, L.K., Ross, L.S., 2001. Apoptosis in the Developing Zebrafish Embryo. Developmental Biology 240, 123–142. 10.1006/dbio.2001.0432

Dekkers, M.P.J., Nikoletopoulou, V., Barde, Y.-A., 2013. Death of developing neurons: New insights and implications for connectivity. Journal of Cell Biology 203, 385–393. 10.1083/jcb.201306136

Dhaka, A., Viswanath, V., Patapoutian, A., 2006. TRP ION CHANNELS AND TEMPERATURE SENSATION. Annu. Rev. Neurosci. 29, 135–161. 10.1146/annurev.neuro.29.051605.112958

Dong, H.P., Holth, A., Ruud, M.G., Emilsen, E., Risberg, B., Davidson, B., 2011. Measurement of apoptosis in cytological specimens by flow cytometry: comparison of Annexin V, caspase cleavage and dUTP incorporation assays. Cytopathology 22, 365–372. 10.1111/j.1365-2303.2010.00811.x

Douglass, A.D., Kraves, S., Deisseroth, K., Schier, A.F., Engert, F., 2008. Escape Behavior Elicited by Single, Channelrhodopsin-2-Evoked Spikes in Zebrafish Somatosensory Neurons. Current Biology 18, 1133–1137. 10.1016/j.cub.2008.06.077

Dyck, P.J., Thomas, P.K., 2005. Peripheral neuropathy, 4th ed. ed. Saunders, Philadelphia.

Eichler, V.B., Porter, R.A., 1981. Rohon-Beard cells in frog development: A study of temporal and spatial changes in a transient cell population. J of Comparative Neurology 203, 121–130. 10.1002/cne.902030110

Eisen, J., 1991. Developmental neurobiology of the zebrafish. J. Neurosci. 11, 311–317. 10.1523/JNEUROSCI.11-02-00311.1991

Eisen, J.S., Pike, S.H., 1991. The spt-1 mutation alters segmental arrangement and axonal development of identified neurons in the spinal cord of the embryonic zebrafish. Neuron 6, 767–776. 10.1016/0896-6273(91)90173-W

Fontenas, L., Kucenas, S., 2021. Spinal cord precursors utilize neural crest cell mechanisms to generate hybrid peripheral myelinating glia. eLife 10, e64267. 10.7554/eLife.64267

Freud, S., 1878. Über Spinalganglien und Rückenmark des Petromyzon.

Freud, S., 1877. Über den Ursprung der hinteren Nervenwurzeln im Rückenmark von Ammocoetes (Petromyzon planeri) Sber. preuss. Altad. Wiss. 75, 15–30.

Frotscher, M., 1997. Dual role of Cajal-Retzius cells and reelin in cortical development. Cell and Tissue Research 290, 315–322. 10.1007/s004410050936

Grunwald, D.J., Kimmel, C.B., Westerfield, M., Walker, C., Streisinger, G., 1988. A neural degeneration mutation that spares primary neurons in the zebrafish. Developmental Biology 126, 115–128. 10.1016/0012-1606(88)90245-X

Ham, T.J., Mapes, J., Kokel, D., Peterson, R.T., 2010. Live imaging of apoptotic cells in zebrafish. FASEB j. 24, 4336–4342. 10.1096/fj.10-161018

Harris, J.E., Whiting, H.P., 1954. Structure and Function in the Locomotory System of the Dogfish Embryo. the Myogenic Stage of Movement. Journal of Experimental Biology 31, 501–524. 10.1242/jeb.31.4.501

Hartenstein, V., 1993. Early pattern of neuronal differentiation in the *Xenopus* embryonic brainstem and spinal cord. J of Comparative Neurology 328, 213–231. 10.1002/cne.903280205

Henderson, K.W., Menelaou, E., Hale, M.E., 2019. Sensory neurons in the spinal cord of zebrafish and their local connectivity. Current Opinion in Physiology 8, 136–140. 10.1016/j.cophys.2019.01.008

Henderson, K.W., Roche, A., Menelaou, E., Hale, M.E., 2020. Hindbrain and Spinal Cord Contributions to the Cutaneous Sensory Innervation of the Larval Zebrafish Pectoral Fin. Front. Neuroanat. 14, 581821. 10.3389/fnana.2020.581821

Henion, P.D., Raible, D.W., Beattie, C.E., Stoesser, K.L., Weston, J.A., Eisen, J.S., 1996. Screen for mutations affecting development of zebrafish neural crest. Dev. Genet. 18, 11–17. 10.1002/(SICI)1520-6408(1996)18:1<11::AID-DVG2>3.0.CO;2-4

Hidalgo, A., Brand, A.H., 1997. Targeted neuronal ablation: the role of pioneer neurons in guidance and fasciculation in the CNS of *Drosophila*. Development 124, 3253–3262. 10.1242/dev.124.17.3253

Hirata, H., Iida, A. (Eds.), 2018. Zebrafish, Medaka, and Other Small Fishes: New Model Animals in Biology, Medicine, and Beyond, 1st ed. 2018. ed. Springer Singapore : Imprint: Springer, Singapore. 10.1007/978-981-13-1879-5

Honjo, Y., Payne, L., Eisen, J.S., 2011. Somatosensory mechanisms in zebrafish lacking dorsal root ganglia: Somatosensory mechanisms in juvenile zebrafish. Journal of Anatomy 218, 271–276. 10.1111/j.1469-7580.2010.01337.x

Hubbard, J.M., Böhm, U.L., Prendergast, A., Tseng, P.-E.B., Newman, M., Stokes, C., Wyart, C., 2016. Intraspinal Sensory Neurons Provide Powerful Inhibition to Motor Circuits Ensuring Postural Control during Locomotion. Current Biology 26, 2841–2853. 10.1016/j.cub.2016.08.026

Hughes, A., 1957. The development of the primary sensory system in Xenopus laevis (Daudin). J Anat 91, 323–338.

Jacobson, M., 1981. Rohon-Beard neurons arise from a substitute ancestral cell after removal of the cell from which they normally arise in the 16-cell frog embryo. J. Neurosci. 1, 923–927. 10.1523/JNEUROSCI.01-08-00923.1981

Joya, X., Garcia-Algar, O., Vall, O., Pujades, C., 2014. Transient Exposure to Ethanol during Zebrafish Embryogenesis Results in Defects in Neuronal Differentiation: An Alternative Model System to Study FASD. PLoS ONE 9, e112851. 10.1371/journal.pone.0112851

Kaji, T., Artinger, K.B., 2004. dlx3b and dlx4b function in the development of Rohon-Beard sensory neurons and trigeminal placode in the zebrafish neurula. Developmental Biology 276, 523–540. 10.1016/j.ydbio.2004.09.020

Kanungo, J., Li, B.-S., Zheng, Y., Pant, H.C., 2006. Cyclin-dependent kinase 5 influences Rohon-Beard neuron survival in zebrafish. J Neurochem 99, 251–259. 10.1111/j.1471-4159.2006.04114.x

Kellermeyer, R., Heydman, L., Mastick, G., Kidd, T., 2018. The Role of Apoptotic Signaling in Axon Guidance. JDB 6, 24. 10.3390/jdb6040024

Kerr, J.F.R., Wyllie, A.H., Currie, A.R., 1972. Apoptosis: A Basic Biological Phenomenon with Wideranging Implications in Tissue Kinetics. Br J Cancer 26, 239–257. 10.1038/bjc.1972.33

Kimmel, C.B., Ballard, W.W., Kimmel, S.R., Ullmann, B., Schilling, T.F., 1995. Stages of embryonic development of the zebrafish. Developmental Dynamics 203, 253–310. 10.1002/aja.1002030302

Kimmel, C.B., Westerfield, M., 1990. Primary neurons of the zebrafish. Signals and sense 561–588.

Knafo, S., Fidelin, K., Prendergast, A., Tseng, P.-E.B., Parrin, A., Dickey, C., Böhm, U.L., Figueiredo, S.N., Thouvenin, O., Pascal-Moussellard, H., Wyart, C., 2017. Mechanosensory neurons control the timing of spinal microcircuit selection during locomotion. eLife 6, e25260. 10.7554/eLife.25260

Kollros, J.J., Bovbjerg, A.M., 1997. Growth and death of Rohon-Beard cells inRana pipiens andCeratophrys ornata. J. Morphol. 232, 67–78. 10.1002/(SICI)1097-4687(199704)232:1<67::AID-JMOR4>3.0.CO;2-L

Kucenas, S., Soto, F., Cox, J.A., Voigt, M.M., 2006. Selective labeling of central and peripheral sensory neurons in the developing zebrafish using P2X(3) receptor subunit transgenes. Neuroscience 138, 641–652. 10.1016/j.neuroscience.2005.11.058

Kwan, K.M., Otsuna, H., Kidokoro, H., Carney, K.R., Saijoh, Y., Chien, C.-B., 2012. A complex choreography of cell movements shapes the vertebrate eye. Development 139, 359–372. 10.1242/dev.071407

Lamborghini, J.E., 1987. Disappearance of Rohon-Beard neurons from the spinal cord of larval *Xenopus laevis*. J of Comparative Neurology 264, 47–55. 10.1002/cne.902640105

Le Douarin, N., Kalcheim, C., 1999. The neural crest, 2nd ed. ed, Developmental and cell biology series. Cambridge university press, Cambridge.

Madabhushi, R., Gao, F., Pfenning, A.R., Pan, L., Yamakawa, S., Seo, J., Rueda, R., Phan, T.X., Yamakawa, H., Pao, P.-C., Stott, R.T., Gjoneska, E., Nott, A., Cho, S., Kellis, M., Tsai, L.-H., 2015. Activity-Induced DNA Breaks Govern the Expression of Neuronal Early-Response Genes. Cell 161, 1592–1605. 10.1016/j.cell.2015.05.032

Malin, J.Z., Shaham, S., 2015. Cell Death in C. elegans Development, in: Current Topics in Developmental Biology. Elsevier, pp. 1–42. 10.1016/bs.ctdb.2015.07.018

Martin, S.C., Sandell, J.H., Heinrich, G., 1998. Zebrafish TrkC1 and TrkC2 Receptors Define Two Different Cell Populations in the Nervous System during the Period of Axonogenesis. Developmental Biology 195, 114–130. 10.1006/dbio.1997.8839

Mazaheri, F., Breus, O., Durdu, S., Haas, P., Wittbrodt, J., Gilmour, D., Peri, F., 2014. Distinct roles for BAI1 and TIM-4 in the engulfment of dying neurons by microglia. Nat Commun 5, 4046. 10.1038/ncomms5046

McConnell, S.K., Ghosh, A., Shatz, C.J., 1989. Subplate Neurons Pioneer the First Axon Pathway from the Cerebral Cortex. Science 245, 978–982. 10.1126/science.2475909

Meltzer, S., Santiago, C., Sharma, N., Ginty, D.D., 2021. The cellular and molecular basis of somatosensory neuron development. Neuron 109, 3736–3757. 10.1016/j.neuron.2021.09.004

Metcalfe, W.K., Myers, P.Z., Trevarrowf, B., Basst, M.B., Kimmel, C.B., 1990. Primary neurons that express the L2/HNK-1 carbohydrate during early development in the zebrafish. Development 110, 491–504. 10.1242/dev.110.2.491

Metcalfe, W.K., Westerfield, M., 1990. Primary Motoneurons of the Zebrafish, in: Raymond, P.A., Easter, S.S., Innocenti, G.M. (Eds.), Systems Approaches to Developmental Neurobiology. Springer US, Boston, MA, pp. 41–47. 10.1007/978-1-4684-7281-3_5

Meyer, G., Soria, J.M., Martínez-Galán, J.R., Martín-Clemente, B., Fairén, A., 1998. Different origins and developmental histories of transient neurons in the marginal zone of the fetal and neonatal rat cortex. J. Comp. Neurol. 397, 493–518. 10.1002/(SICI)1096-9861(19980810)397:4<493::AID-CNE4>3.0.CO;2-X

Moreno, R.L., Ribera, A.B., 2014. Spinal neurons require Islet1 for subtype-specific differentiation of electrical excitability. Neural Dev 9, 19. 10.1186/1749-8104-9-19

Mukherjee, A., Williams, D.W., 2017. More alive than dead: non-apoptotic roles for caspases in neuronal development, plasticity and disease. Cell Death Differ 24, 1411–1421. 10.1038/cdd.2017.64

Nakano, Y., Fujita, M., Ogino, K., Saint-Amant, L., Kinoshita, T., Oda, Y., Hirata, H., 2010. Biogenesis of GPI-anchored proteins is essential for surface expression of sodium channels in zebrafish Rohon-Beard neurons to respond to mechanosensory stimulation. Development 137, 1689– 1698. 10.1242/dev.047464

Nieuwenhuys, R., 1964. Comparative Anatomy of the Spinal Cord, in: Progress in Brain Research. Elsevier, pp. 1–57. 10.1016/S0079-6123(08)64043-1

Nishikawa, A., 2012. Cell Interaction During Larval-To-Adult Muscle Remodeling in the Frog, Xenopus laevis, in: Gowder, S. (Ed.), Cell Interaction. InTech. 10.5772/47757

Nordlander, R.H., 1989. HNK-1 marks earliest axonal outgrowth in Xenopus. Developmental Brain Research 50, 147–153. 10.1016/0165-3806(89)90135-1

O’Brien, G.S., Rieger, S., Wang, F., Smolen, G.A., Gonzalez, R.E., Buchanan, J., Sagasti, A., 2012. Coordinate development of skin cells and cutaneous sensory axons in zebrafish. J Comp Neurol 520, 816–831. 10.1002/cne.22791

Ogino, K., Hirata, H., 2018. Rohon-Beard Neuron in Zebrafish, in: Hirata, H., Iida, A. (Eds.), Zebrafish, Medaka, and Other Small Fishes. Springer Singapore, Singapore, pp. 59–81. 10.1007/978-981-13-1879-5_4

Palanca, A.M.S., Lee, S.-L., Yee, L.E., Joe-Wong, C., Trinh, L.A., Hiroyasu, E., Husain, M., Fraser, S.E., Pellegrini, M., Sagasti, A., 2013. New transgenic reporters identify somatosensory neuron subtypes in larval zebrafish. Devel Neurobio 73, 152–167. 10.1002/dneu.22049

Park, B.-Y., Hong, C.-S., Weaver, J.R., Rosocha, E.M., Saint-Jeannet, J.-P., 2012. Xaml1/Runx1 is required for the specification of Rohon-Beard sensory neurons in Xenopus. Developmental Biology 362, 65–75. 10.1016/j.ydbio.2011.11.016

Patten, S.A., Sihra, R.K., Dhami, K.S., Coutts, C.A., Ali, D.W., 2007. Differential expression of PKC isoforms in developing zebrafish. Intl J of Devlp Neuroscience 25, 155–164. 10.1016/j.ijdevneu.2007.02.003

Pineda, R.H., Svoboda, K.R., Wright, M.A., Taylor, A.D., Novak, A.E., Gamse, J.T., Eisen, J.S., Ribera, A.B., 2006. Knockdown of Nav 1.6a Na+ channels affects zebrafish motoneuron development. Development 133, 3827–3836. 10.1242/dev.02559

Pittman, A.J., Law, M.-Y., Chien, C.-B., 2008. Pathfinding in a large vertebrate axon tract: isotypic interactions guide retinotectal axons at multiple choice points. Development 135, 2865– 2871. 10.1242/dev.025049

Pop, S., Chen, C.-L., Sproston, C.J., Kondo, S., Ramdya, P., Williams, D.W., 2020. Extensive and diverse patterns of cell death sculpt neural networks in insects. eLife 9, e59566. 10.7554/eLife.59566

Raible, D.W., Eisen, J.S., 1994. Restriction of neural crest cell fate in the trunk of the embryonic zebrafish. Development 120, 495–503. 10.1242/dev.120.3.495

Raible, D.W., Wood, A., Hodsdon, W., Henion, P.D., Weston, J.A., Eisen, J.S., 1992. Segregation and early dispersal of neural crest cells in the embryonic zebrafish. Dev. Dyn. 195, 29–42. 10.1002/aja.1001950104

Rasmussen, J.P., Vo, N.-T., Sagasti, A., 2018. Fish Scales Dictate the Pattern of Adult Skin Innervation and Vascularization. Developmental Cell 46, 344–359.e4. 10.1016/j.devcel.2018.06.019

Reyes, R., Haendel, M., Grant, D., Melancon, E., Eisen, J.S., 2004. Slow degeneration of zebrafish Rohon-Beard neurons during programmed cell death. Dev. Dyn. 229, 30–41. 10.1002/dvdy.10488

Roberts, A., Clarke, J.D., 1982. The neuroanatomy of an amphibian embryo spinal cord. Philos Trans R Soc Lond B Biol Sci 296, 195–212. 10.1098/rstb.1982.0002

Roberts, A., Smyth, D., 1974. The development of a dual touch sensory system in embryos of the amphibianXenopus laevis. J. Comp. Physiol. 88, 31–42. 10.1007/BF00695921

Rohon, J.V., 1884. Histogenese des Ruckenmarkes der Forelle. Akad Wiss Math.

Rossi, C.C., Hernandez-Lagunas, L., Zhang, C., Choi, I.F., Kwok, L., Klymkowsky, M., Bruk Artinger, K., 2008. Rohon-Beard sensory neurons are induced by BMP4 expressing non-neural ectoderm in Xenopus laevis. Developmental Biology 314, 351–361. 10.1016/j.ydbio.2007.11.036

Rossi, C.C., Kaji, T., Artinger, K.B., 2009. Transcriptional control of Rohon-Beard sensory neuron development at the neural plate border. Dev. Dyn. 238, 931–943. 10.1002/dvdy.21915

Sagasti, A., Guido, M.R., Raible, D.W., Schier, A.F., 2005. Repulsive Interactions Shape the Morphologies and Functional Arrangement of Zebrafish Peripheral Sensory Arbors. Current Biology 15, 804–814. 10.1016/j.cub.2005.03.048

Saint-Amant, L., Drapeau, P., 1998. Time course of the development of motor behaviors in the zebrafish embryo. J. Neurobiol. 37, 622–632. 10.1002/(SICI)1097-4695(199812)37:4<622::AID-NEU10>3.0.CO;2-S

Schindelin, J., Arganda-Carreras, I., Frise, E., Kaynig, V., Longair, M., Pietzsch, T., Preibisch, S., Rueden, C., Saalfeld, S., Schmid, B., Tinevez, J.-Y., White, D.J., Hartenstein, V., Eliceiri, K., Tomancak, P., Cardona, A., 2012. Fiji: an open-source platform for biological-image analysis. Nat Methods 9, 676–682. 10.1038/nmeth.2019

Segawa, H., Miyashita, T., Hirate, Y., Higashijima, S., Chino, N., Uyemura, K., Kikuchi, Y., Okamoto, H., 2001. Functional Repression of Islet-2 by Disruption of Complex with Ldb Impairs Peripheral Axonal Outgrowth in Embryonic Zebrafish. Neuron 30, 423–436. 10.1016/S0896-6273(01)00283-5

Shorey, M., Rao, K., Stone, M.C., Mattie, F.J., Sagasti, A., Rolls, M.M., 2021. Microtubule organization of vertebrate sensory neurons in vivo. Developmental Biology 478, 1–12. 10.1016/j.ydbio.2021.06.007

Spitzer, N.C., 1984. What do rohon-beard cells do? Trends in Neurosciences 7, 224–225. 10.1016/S0166-2236(84)80208-8

Spitzer, N.C., 1982. Voltage- and stage-development uncoupling of Rohon-Beard neurones during embryonic development of *Xenopus* tadpoles. The Journal of Physiology 330, 145–162. 10.1113/jphysiol.1982.sp014334

Suster, M.L., Kikuta, H., Urasaki, A., Asakawa, K., Kawakami, K., 2009. Transgenesis in Zebrafish with the Tol2 Transposon System, in: Cartwright, E.J. (Ed.), Transgenesis Techniques, Methods in Molecular Biology. Humana Press, Totowa, NJ, pp. 41–63. 10.1007/978-1-60327-019-9_3

Svoboda, K.R., Linares, A.E., Ribera, A.B., 2001. Activity regulates programmed cell death of zebrafish Rohon-Beard neurons. Development 128, 3511–3520. 10.1242/dev.128.18.3511

Takamiya, M., Campos-Ortega, J.A., 2006. Hedgehog signalling controls zebrafish neural keel morphogenesis via its level-dependent effects on neurogenesis. Developmental Dynamics 235, 978–997. 10.1002/dvdy.20720

Tamme, R., Wells, S., Conran, J.G., Lardelli, M., 2002. The identity and distribution of neural cells expressing the mesodermal determinant spadetail. BMC Dev Biol 2, 9. 10.1186/1471-213X-2-9

Theveneau, E., Mayor, R., 2012. Neural crest delamination and migration: From epithelium-to-mesenchyme transition to collective cell migration. Developmental Biology 366, 34–54. 10.1016/j.ydbio.2011.12.041

Tongiorgi, E., Bernhardt, R.R., Schachner, M., 1995. Zebrafish neurons express two L1-related molecules during early axonogenesis. J of Neuroscience Research 42, 547–561. 10.1002/jnr.490420413

Tuttle, A.M., Miller, L.N., Royer, L.J., Wen, H., Kelly, J.J., Calistri, N.L., Heiser, L.M., Nechiporuk, A.V., 2024. Single-cell analysis of Rohon-Beard neurons implicates Fgf signaling in axon maintenance and cell survival. J. Neurosci. e1600232024. 10.1523/JNEUROSCI.1600-23.2024

Uemura, O., Okada, Y., Ando, H., Guedj, M., Higashijima, S., Shimazaki, T., Chino, N., Okano, H., Okamoto, H., 2005. Comparative functional genomics revealed conservation and diversification of three enhancers of the isl1 gene for motor and sensory neuron-specific expression. Developmental Biology 278, 587–606. 10.1016/j.ydbio.2004.11.031

Umeda, K., Ishizuka, T., Yawo, H., Shoji, W., 2016. Position- and quantity-dependent responses in zebrafish turning behavior. Sci Rep 6, 27888. 10.1038/srep27888

Venero Galanternik, M., Navajas Acedo, J., Romero-Carvajal, A., Piotrowski, T., 2016. Imaging collective cell migration and hair cell regeneration in the sensory lateral line, in: Methods in Cell Biology. Elsevier, pp. 211–256. 10.1016/bs.mcb.2016.01.004

Whitlock, K.E., Westerfield, M., 1998. A Transient Population of Neurons Pioneers the Olfactory Pathway in the Zebrafish. J. Neurosci. 18, 8919–8927. 10.1523/JNEUROSCI.18-21-08919.1998

Williams, J.A., Barrios, A., Gatchalian, C., Rubin, L., Wilson, S.W., Holder, N., 2000. Programmed Cell Death in Zebrafish Rohon Beard Neurons Is Influenced by TrkC1/NT-3 Signaling. Developmental Biology 226, 220–230. 10.1006/dbio.2000.9860

Williams, K., Ribera, A.B., 2020. Long-lived zebrafish Rohon-Beard cells. Developmental Biology 464, 45–52. 10.1016/j.ydbio.2020.05.003

Wilson, S.W., Easter, S.S., 1991. A pioneering growth cone in the embryonic zebrafish brain. Proc. Natl. Acad. Sci. U.S.A. 88, 2293–2296. 10.1073/pnas.88.6.2293

Won, Y.-J., Ono, F., Ikeda, S.R., 2012. Characterization of Na+ and Ca2+ channels in zebrafish dorsal root ganglion neurons. PLoS One 7, e42602. 10.1371/journal.pone.0042602

Won, Y.-J., Ono, F., Ikeda, S.R., 2011. Identification and Modulation of Voltage-Gated Ca ^2+^ Currents in Zebrafish Rohon-Beard Neurons. Journal of Neurophysiology 105, 442–453. 10.1152/jn.00625.2010

Woolf, C.J., Ma, Q., 2007. Nociceptors—Noxious Stimulus Detectors. Neuron 55, 353–364. 10.1016/j.neuron.2007.07.016

Wright, M.A., Ribera, A.B., 2010. Brain-Derived Neurotrophic Factor Mediates Non-Cell-Autonomous Regulation of Sensory Neuron Position and Identity. J. Neurosci. 30, 14513– 14521. 10.1523/JNEUROSCI.4025-10.2010

